# An image-based biophysical model of the lung to investigate the effect of pulmonary surfactant on lung function

**DOI:** 10.1101/2025.02.15.638361

**Authors:** Sunder Neelakantan, Kyle J. Myers, Rahim Rizi, Bradford J. Smith, Reza Avazmohammadi

## Abstract

Lung biomechanics aims to understand the structure-function relationship in the lung under normal and pathological conditions. Pulmonary surfactant has a key structural role in lung function with a significant contribution to the mechanical response of the lungs during respiration. Pulmonary surfactant dynamically regulates surface tension at the air-liquid interface to decrease stiffness during respiration and prevent alveolar collapse during lower volumes. Many lung injuries involve alterations to the contribution of surfactants to lung function. We developed a novel biophysical model using a poroelastic formulation that incorporates pulmonary surfactant dynamics and aims to quantify the contribution of the pulmonary surfactant toward lung compliance. The effect of pulmonary surfactant was modeled as a surface energy function, and the surface behavior was converted to bulk behavior by assuming uniform spherical alveoli. The model was used to simulate respiration and investigate the effect of altered surface tension caused by surfactant dysfunction. The model captured the characteristic sigmoidal inspiratory pressure-volume curve and hysteresis observed during clinical measurements. In addition, the model predicted the expected behavior in surfactant dysfunction in lung injuries. We expect this work to serve as an essential step towards de-convoluting and predicting the contributions of the lung parenchyma and pulmonary surfactant to global and regional lung compliance in health and disease.

## 1 Introduction

Lung injuries and diseases such as acute respiratory disease syndrome (ARDS), emphysema, and ventilator-induced lung injuries (VILI) can cause significant alterations to the biomechanical behavior of the lung parenchyma. These alterations in lung biomechanics can impair lung function and lead to poor oxygenation and carbon dioxide clearance. In addition, the mechanical stresses of respiration modulate the lung function at a cellular level through mechanotransduction^1, 2^. An essential component of this mechanical behavior is the effect of pulmonary surfactant, which is present in the thin liquid film that covers the alveolar septal walls. Many pathological conditions, including ARDS^3–5^ and premature birth^6, 7^ adversely affect the pulmonary surfactant system, leading to increased surface tension, atelectasis, and a reduction of gas exchanging surface area (Fig. 1). This alteration in mechanical response is spatially heterogeneous and, as such, can lead to ventilation heterogeneities. The ventilation heterogeneities can significantly impact the efficacy of one of the most common clinical intervention strategies - mechanical ventilation and can lead to further damage known as ventilator-induced lung injuries (VILI)^8^.

**Fig. 1.**
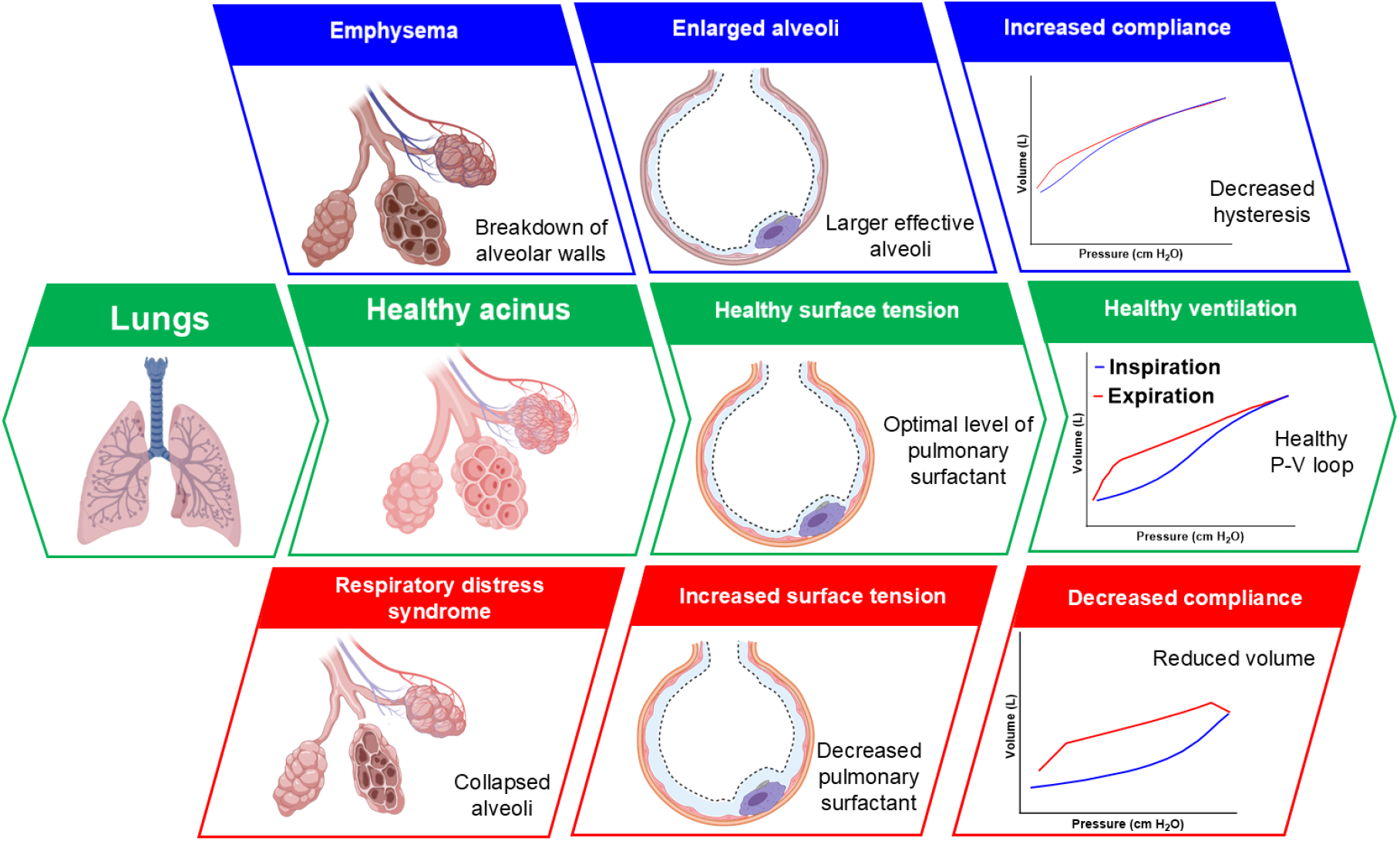
Effect of altered surface tension and initial alveolar radius on lung ventilation. These alterations lead to substantial changes in lung compliance and, subsequently, lung volume.

To understand pathologic lung function due to alterations in pulmonary surfactant regulation and/or alterations in parenchymal elasticity, it proves necessary to first understand the mechanical response of the healthy lung. The two major primary drivers of lung mechanical function are the viscoelastic properties of the parenchymal tissue and the surface tension of the liquid film containing the pulmonary surfactant. While the ex-vivo tissue elasticity of lung parenchyma has been extensively studied and characterized^9–11^, the ex-vivo characterization of pulmonary surfactant mechanics remains challenging. Studies have investigated the surface tension of pulmonary surfactant through in-situ^12, 13^ and in-vitro methods^14–16^. However, evaluating the effect of pulmonary surfactant as part of parenchymal tissue ex vivo remains understudied, concerning two limitations. The first limitation is that pulmonary surfactant breaks down in the alveoli and is replenished by alveolar epithelial Type II cells^17^. When parenchymal tissue is excised, the surfactant continues to break down but is not replenished due to the absence of functioning epithelial cells. The second limitation of ex-vivo testing is that parenchymal tissue specimens need to be submerged to remain viable during testing, which can lead to the surfactant being washed away.

Computational lung modeling is one of the methods to investigate lung biomechanics^18–20^ that has gained increasing interest recently due to its potential to provide tools to investigate various ventilation scenarios prior to clinical implementation^21^. The simulation of ventilation-induced stress and strain distributions allows for optimizing ventilator settings to provide a non-damaging micro-mechanical environment in the lung^22^. Studies have incorporated the effect of pulmonary surfactant into reduced order models^23, 24^ and into acinar models of the lungs^25, 26^. Many lung models have also incorporated the effect of varying surface tension^25, 27–31^ to mimic the observed hysteresis in the lung pressure-volume relationship. Among others, Wiechert et al.^25^ developed an alveolar model that accounted for the varying surface tension of the liquid film. They modeled a single alveolar sac as a truncated octahedron and incorporated the dynamic surface tension of the liquid film by including an interfacial energy term on the inner surface of the tetrakaidecahedron (TKD) (alveolar) element. Another study, by Ma et al.^31^, incorporated the effect of pulmonary surfactant by modeling the alveoli as truncated octahedrons^28–30^ and accounted for surfactant transport during the respiration cycle. This model allowed for the adjustment of surface tension inside each alveolus and successfully recapitulated the pressure-volume behavior observed during measurements^32^. However, the primary limitation of such models was the need to model individual alveoli using polyhedral shapes.

Modeling pulmonary surfactant dynamics in a whole lung remains a challenging research gap due to the tremendous disparity in length scales, thin-film interfacial flows, surfactant physicochemical hydrodynamics, and fluid-structure interactions in the lung. To address this challenge, we developed a model incorporating pulmonary surfactant effects into a continuum-level mechanical model of the lungs. To reduce the computational costs and enhance the translational potential of the model for clinical applications, we utilized a continuum poroelastic model to represent the bulk parenchymal tissue and invoked the concept of surface energy to incorporate dynamic surface tension caused by the pulmonary surfactant. We used our model to investigate the effect of various physiological parameters, such as alveoli’s density and radius and surface tension’s strength, on the organ-level lung mechanical function. Our model recapitulates the organ-level behavior of the lungs in various diseases that affect the alveolar surface area and the regulation of pulmonary surfactant such as ARDS and VILI.

## 2 Methods

### 2.1 Constitutive modeling of lung parenchyma and pulmonary surfactant

We modeled the lung as a poroelastic material representing porous parenchymal tissue consisting of soft tissue skeleton, air, and the air-liquid interface at the surface of the parenchymal tissue where pulmonary surfactant dynamically alters surface tension. The bulk soft tissue was assumed to be a compressible neo-Hookean hyperelastic material whose strain energy contribution (*U*^*B*^) is given by

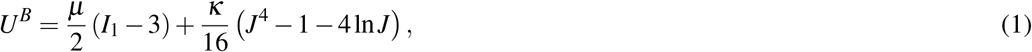

where *µ* is the shear modulus and *κ* is the bulk modulus. *I*_1_ and *I*_3_ = *J*^2^ are the first and third invariants of the Cauchy green strain tensor (**C**). **C** is calculated using the deformation gradient tensor (**F**) as **C** = **F**^*T*^ **F**, where *F*_*i j*_ = *δ*_*i j*_ + *∂u*_*i*_*/∂X*_*j*_. Here *u*_*i*_ and *X*_*j*_ are the displacement and position vectors in the reference frame, and **I** is the identity tensor. **C** was used to define the strain tensor **E**, given by **E** = (**C** − **I**)*/*2.

The strain energy contribution of the liquid film surface tension is given by the *U*^*S*^ as a function of surface tension and surface area ratio^33, 34^, written as

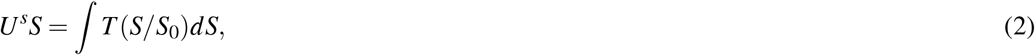

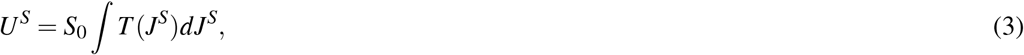

where *T* (*S/S*_0_) is the surface tension of the liquid film that is a function of the surface area ratio *S/S*_0_. The variation of *T* with surface area ratio was adapted from the in-vitro study by Otis et al.^14^. *S* and *S*_0_ are the total alveolar surface area in the current and reference configurations, with the latter corresponding to the end-expiration timepoint. 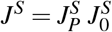 is the total surface area ratio where *J*^*S*^ is the change in surface area from reference to the current configuration, and 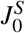 is the change in surface area from a hypothetical stress-free configuration (corresponding to a zero pressure state) to the reference configuration.

Next, Eqs. 2,3 were converted from surface terms to volumetric (bulk) terms such that the surface energy contributions of the parenchymal tissue and surfactant can be summed. We considered a representative tissue volume cube (*V*_0_ = 1m^3^) of parenchymal tissue. The porosity of a given volume was taken to be *ϕ*, which is the fraction of air volume to total volume. We assume this volume contains *n* perfectly spherical alveoli with a uniform initial radius *r*_0_ such that

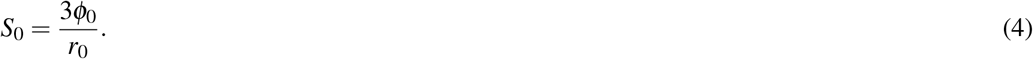

The cumulative work done to increase the surface area ratio is represented by *F*(*s*), i.e., *F*(*J*^*S*^) = *T* (*J*^*S*^)*dJ*^*S*^. Using Eq. 4, the surface invariant *J*^*S*^ can be expressed in terms of the bulk invariant *J*, given by

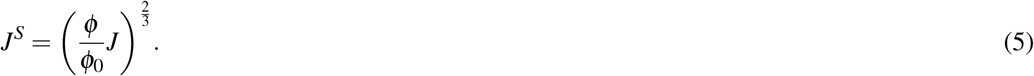

Now, given the constant volume of the representative tissue cube, we have

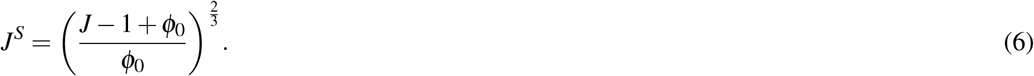

Thus, the surface energy function can be expressed in terms of structural constants and deformation invariants, given by

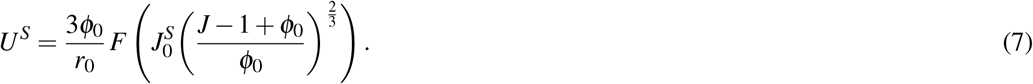

The variation of surface tension *T* (*J*^*S*^) as a function of the total surface area ratio *J*^*S*^ was adopted from an in-vitro study by Otis et al.^14^ shown in Fig. 2C. Combining the bulk and surface contributions, the net strain energy function (*U*) was obtained as

**Fig. 2.**
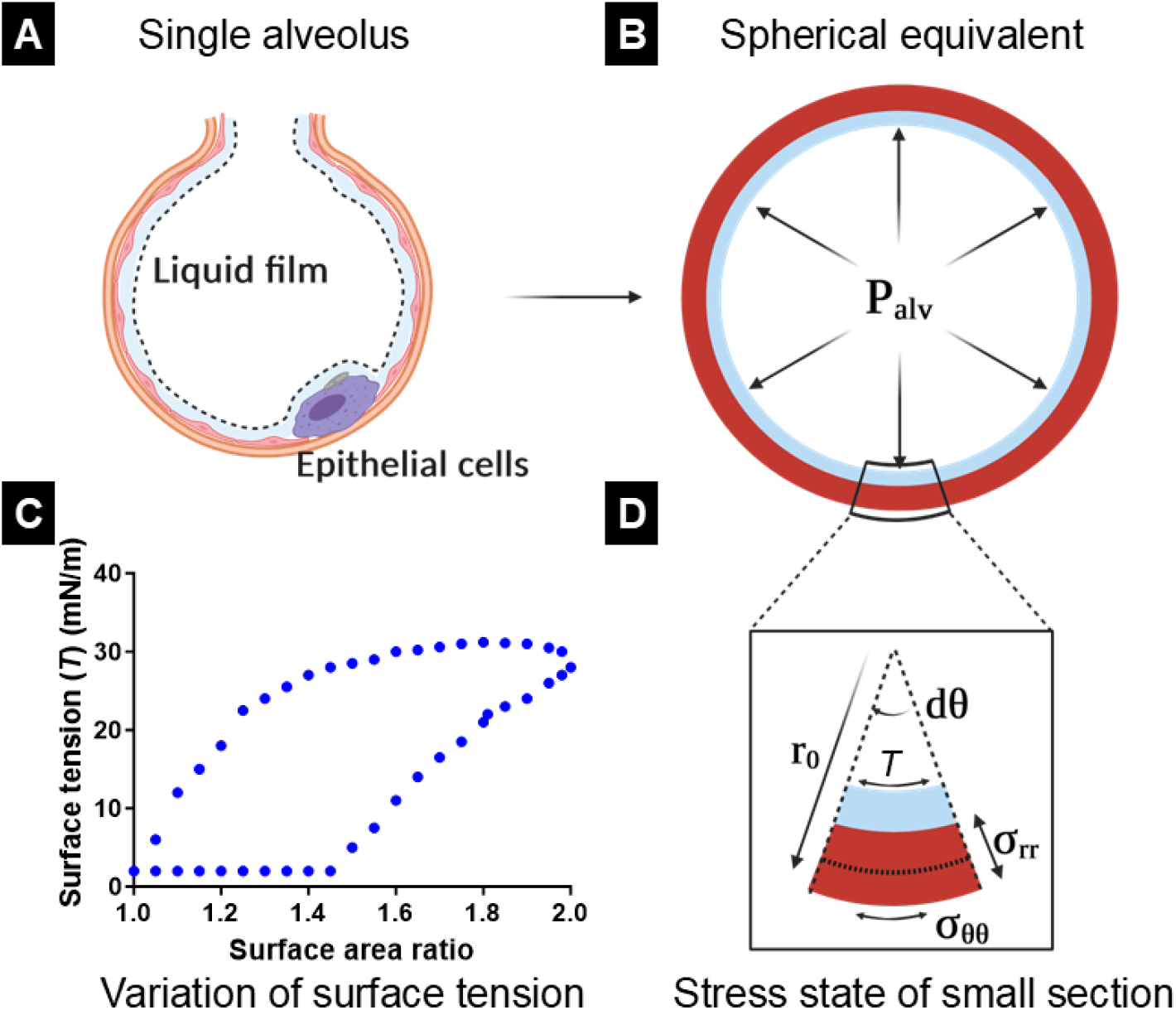
(**A**) The structure of a single alveolus, (**B**) the mechanical equivalent used in the model, (**C**) the variation of surface tension as a function of alveolar surface area deformation (adapted from Otis et al.^14^) and (**D**) A small section of the alveoli to describe the stresses acting on the “alveolar sphere”. *P*_*alv*_ - alveolar pressure, *r*_0_ - initial radius of alveoli, *dθ* - small angle, *T* - effect of surface tension on small alveolar section, *σ*_*rr*_ - radial stress on alveolar wall, *σ*_*θθ*_ - circumferential stress on alveolar wall.

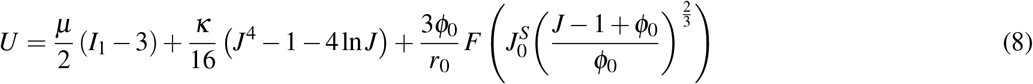

### 2.2 Segmentation and reconstruction of lung geometry

The constitutive modeling in Section 2.1 was implemented in an image-based lung model to deconvolute bulk and surface effects on organ-level lung behavior. The left lung and central airways were segmented from human CT scans (n=1, male anonymous patient scan from the EMPIRE10 repository^35^) using Materialize Mimics (Fig. 3). The segmented left lung was reconstructed using 3-matic (Fig. 3). The segmented airways were used to define the geometry of the airway tree up to the third generation. The airway tree was synthetically extended downstream using a fractal tree algorithm adopted from Tawhai et al.^36^. Briefly, the center of mass of the lung was found and connected to the point of origin of the main bronchus. The airway was generated along this vector direction, with the length of the airway being 0.3 times the distance between the two points. Next, the lung was split into two parts with roughly equal volume, and the process was repeated with each half of the lung till the desired number of airways generations were reached or till the airways reached the surface of the lungs.

**Fig. 3.**
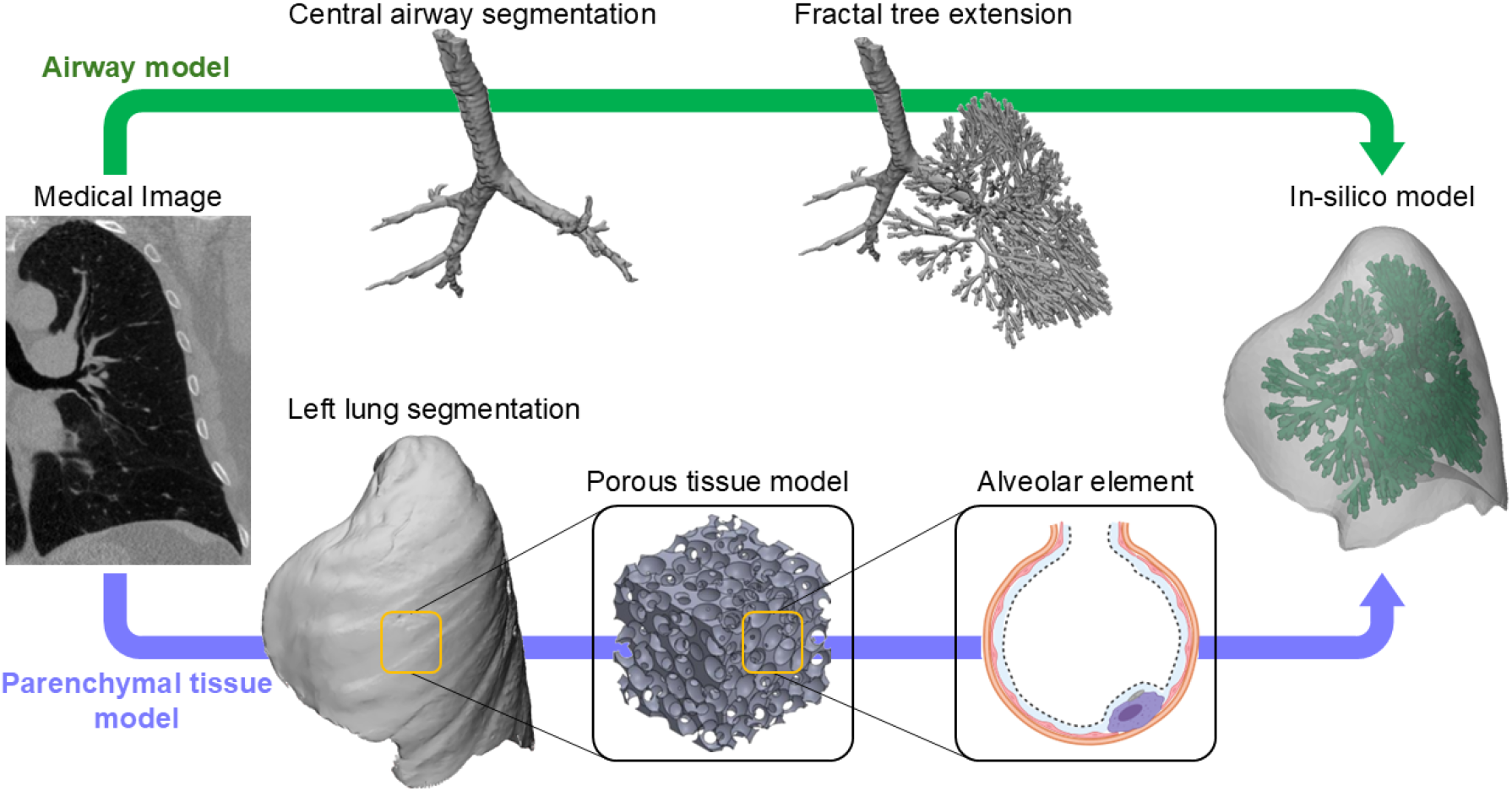
Segmentation and reconstruction of left lung geometry from CT scans. Central airways and the left lung were segmented from CT scans. The central airway was extended using a synthetic tree growth algorithm, and the result was combined with the left lung geometry.

The final meshing process was carried out using Materialize 3-matic. The lumen and the airway walls were removed from the left lung such that the mesh represented only the porous parenchymal tissue. Removing airways exposed the surface area for the fluid to diffuse into the porous parenchyma. In a physiological setting, air diffuses from the airways to the alveoli only from the distal vessels, but this was simplified in our model to improve simulation stability. The acinar units of the lungs are ventilated from the airways and air diffuses between neighboring alveoli through the pores of Kohn to equalize the pressure between multiple alveoli.

### 2.3 In-silico model of the lungs

The simulations of the left lung were carried out in ABAQUS. Time-dependent fluid (air) pressure (as a linear ramp) was applied as input on the entire luminal surface of the airways interfacing with the parenchymal tissues. The maximum pressure was set to 20 cm H_2_O. At the end-inspiration timepoint, the applied pressure was linearly decreased to simulate expiration. Increases in the pressure lead to fluid (air) perfusion in the lungs, which simulates mechanical ventilation. The regions engulfing airways were held fixed, and spring boundary conditions were applied to the lung surface to simulate the effect of the ribs and diaphragm motion. The change in lung volume was computed as a function of the applied air pressure. The constitutive model parameters of parenchymal elasticity were constant for all the in-silico simulations, with *µ* = 2.92 kPa and *κ* = 0.25 kPa.

#### 2.3.1 Simulating pathologies connecting surface tension and porosity alterations to lung dysfunction

After optimizing the material parameters such that the P-V loop resembled that reported in literature^32^, *U*^*S*^ was altered to investigate the effect of surfactant dysfunction. *U*^*S*^ was scaled to 0, 0.5, 1.5, and 2 times the values of that in the healthy lung to simulate the effect of both increased and decreased surface tension in the alveoli. In addition, the increased initial alveolar radius was also investigated to simulate the effect of emphysema on the P-V loop.

## 3 Results

### 3.1 In-silico model captured hysteresis observed during respiration

The model with the presence of the liquid film and pulmonary surfactant demonstrates hysteresis where, at the same volume, pressures are higher on inspiration than on expiration (Fig. 4A). The inspiration curve also displayed the characteristic sigmoidal shape during the inflation of lungs containing pulmonary surfactant. This qualitative behavior is in agreement with the curves reported in the literature (Fig. 4B)^32^.

**Fig. 4.**
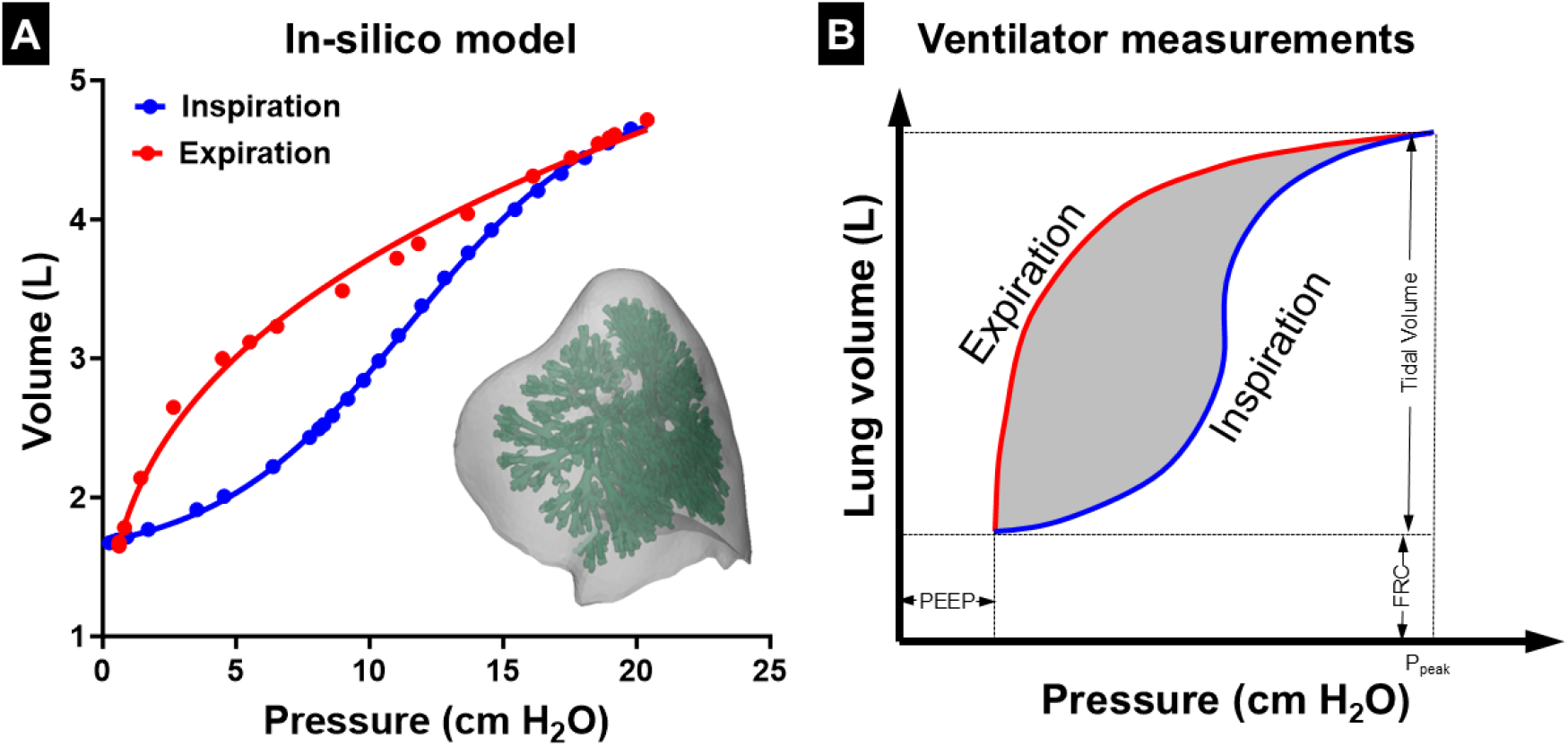
(**A**) Simulation of the lung parenchyma with pulmonary surfactant (neo-Hookean model incorporating surface energy from pulmonary surfactant). Points represent steps in in-silico simulation, and lines are from fitting an appropriate curve to the points. (**B**) Representative respiratory pressure-volume loop measurement adapted from literature^32, 37^. FRC: functional residual capacity, PEEP: positive end expiratory pressure.

### 3.2 Effect of simulated pathology on ventilation

Decreasing the surface tension contribution resulted in reduced hysteresis (Figs. 5A, B). The inflection point was moved to lower pressures as compared to the healthy P-V loop (Figs. 5C). The inflection point moved from 15 cm H_2_O to 10 cm H_2_O when the surface tension was scaled to half the value in the healthy lung (Figs. 5B, C). Moreover, decreasing surface tension to zero caused the P-V loops to shift from a sigmoidal shape (Fig. 5A) to a concave shape, and the curves behaved similarly to a neo-Hookean material (Figs. 5D). In contrast, increased surface tension contribution resulted in the inflection point being moved to higher pressures (Figs. 5C), with the inflection point being greater than 20 cm H_2_O when the surface tension was twice the healthy value. A significant increase in surface tension resulted in the P-V loops turning convex (Figs. 5E). In addition to the qualitative behavior of the P-V loops, increased surface tension also resulted in a significantly lower maximum volume at the same applied pressure, whereas decreased surface tension caused an increase in the maximum volume. These results indicate that increased surface tension substantially reduces lung compliance, as expected.

**Fig. 5.**
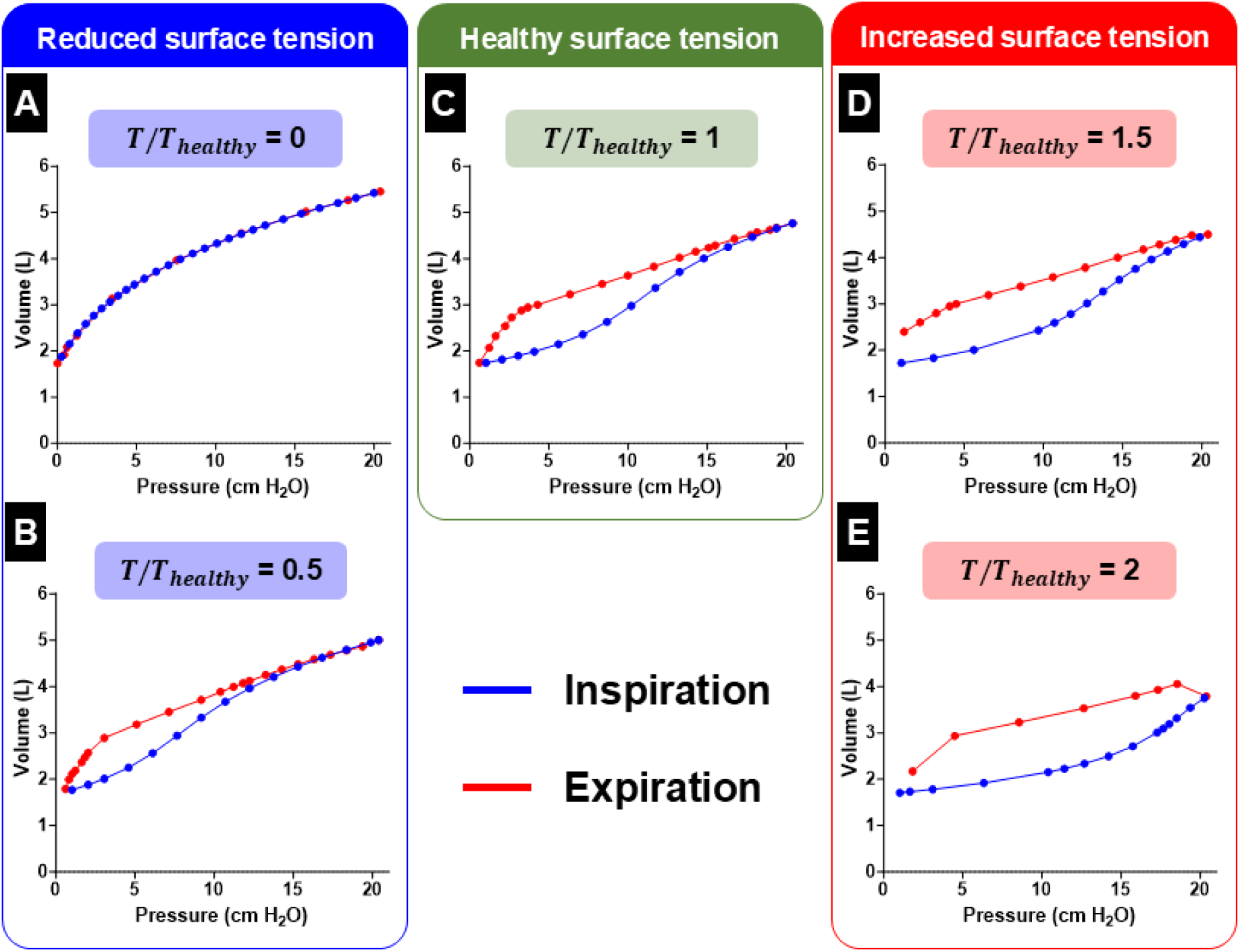
Effect of altered surface tension. (**A**) Zero contribution from surface tension (*T* scaling - 0). (**B**) Mild decrease in surface tension (*T* scaling - 0.5). (**C**) Healthy surface tension (*T* scaling - 1). (**D**) Mild increase in surface tension (*T* scaling - 1.5). (**E**) Strong increase in surface tension (*T* scaling - 2). *T* scaling - Multiplicative factor for the strain energy contribution from the surface tension of the air-to-liquid interface.

In addition, varying the initial radius to simulate the effect of diseases like emphysema indicated significant changes to the behavior of the P-V loop. The hysteresis behavior was substantially diminished (Fig. 6), with the inspiratory and expiratory phases of the P-V loop overlapping with each other. In addition, there was a mild increase in lung compliance, with maximum volume increasing by 7.3% and 9.7%, respectively, when the initial alveolar radius increased from 100*µ*m to 200*µ*m and 300*µ*m.

**Fig. 6.**
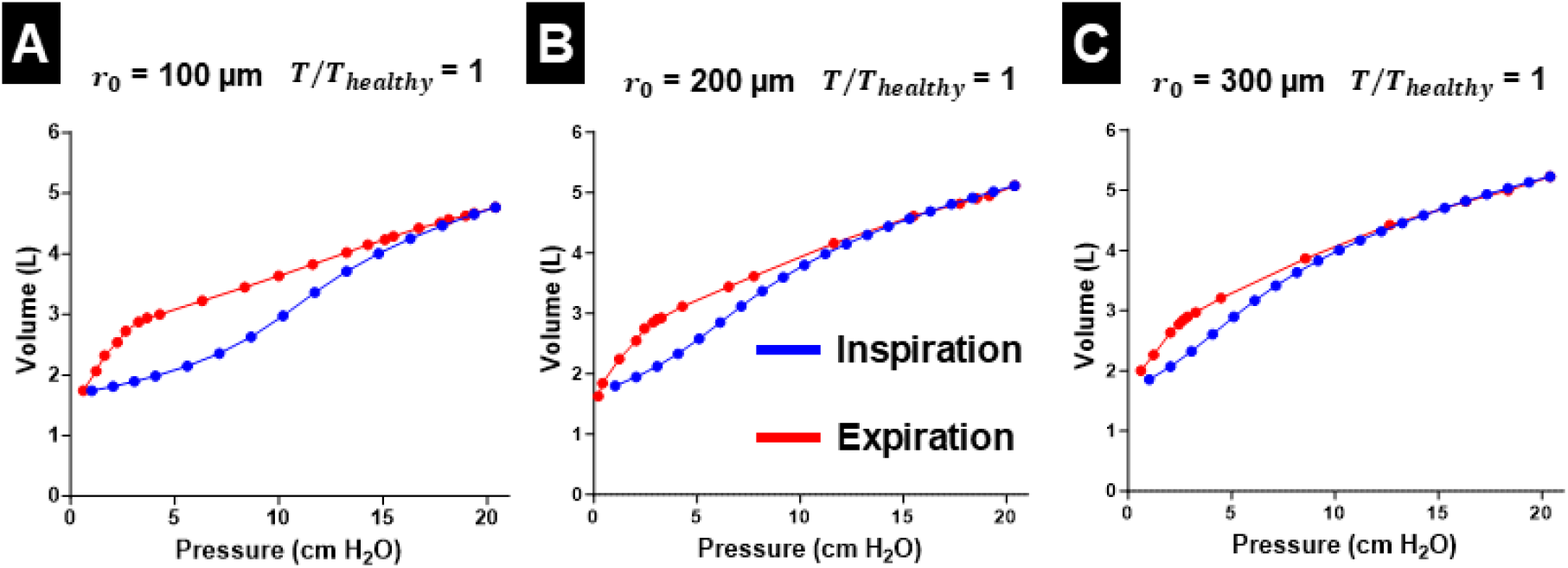
Effect of altered initial alveolar radius (*r*_0_). (**A**) *r*_0_=100*µ*m. (**B**) *r*_0_=200*µ*m. (**C**) *r*_0_=300*µ*m. The surface tension remained constant and set to the same values as the healthy simulation.

### 3.3 Regional kinematic behavior during ventilation

The maximum principal strain (**E**) for the representative simulation is presented in Fig. 7. Strains visualized on the slices show large strains near the fixed airway (Fig. 7A), with decreasing strain closer to the pleural surface (Fig. 7B). The displacement in this representative simulation indicated that the top of the lungs experienced minimal motion, while the region near the diaphragm experienced significant downward displacement (Fig. 7C). The displacement contours from the in-silico model were both qualitatively and quantitatively similar to those reported in the study by Castillo et al.^38^ (Fig. 7D). The motion reported in that study was obtained via dynamic CT imaging using manual displacement measurement at 1166 landmarks. An interesting observation is that the strain contours were different from the displacement contour, and this could be attributed to the inflation of the lung contributing more to the deformation and only minimally to the displacement of the lung tissue.

**Fig. 7.**
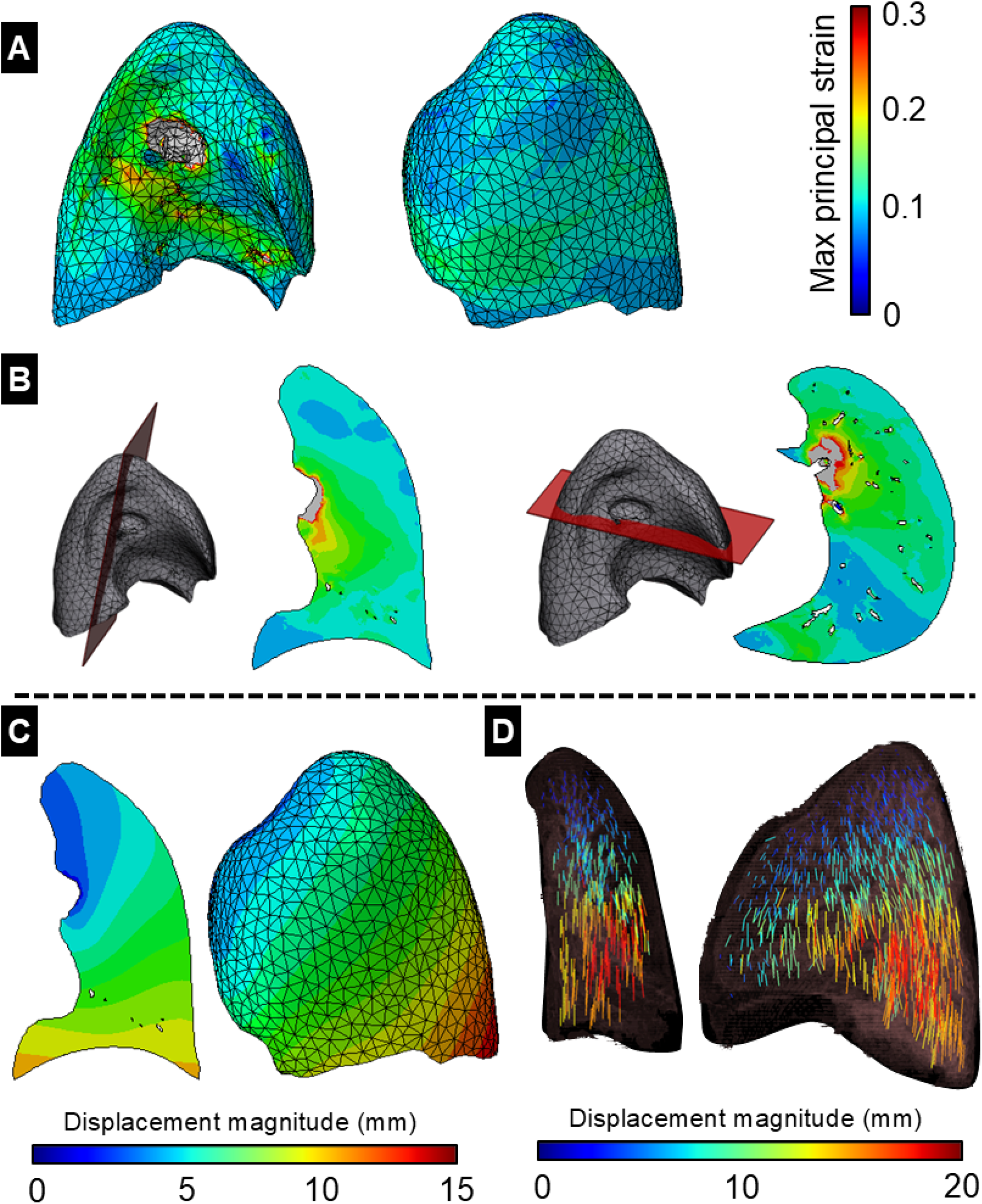
Maximum principal strain at the end of inspiration estimated as the largest eigenvalue of the strain tensor (**E**). Maximum principal strain presented on (**A**) the surface of the lungs, (**B**) slices in the frontal plane and the transverse plane. (**C**) Displacement magnitude along a frontal slice and the surface of the lungs. (**D**) Displacement measured at 1166 landmarks through dynamic CT imaging in the study by Castillo et al.^38^.

## 4 Discussion

### 4.1 In-silico model deconvoluted the contributions of pulmonary surfactant and parenchymal tissue elasticity

In this study, we presented a phenomenological biophysics-based model that incorporates the effects of pulmonary surfactant into a compressible hyperelastic model of lung parenchyma. Using this model, we qualitatively recapitulated the pressure-volume curve in literature^32^. The simulation, including pulmonary surfactant, demonstrated the expected counterclockwise hysteresis with substantially higher stiffness on inspiration than expiration. This hysteresis behavior vanishes when simulating saline inflation, which is expected of a neo-Hookean material model and is in agreement with existing data on ventilation of the lung with saline, which eliminates surface tension^39–41^. The model can also vary physical parameters such as initial porosity (*ϕ*_0_), initial alveolar radius (*r*_0_), and the variation of surface tension with surface deformation in addition to varying the elastic behavior of the lung parenchyma. The capacity to vary these parameters per finite element can enable the investigation of lung diseases and injuries that lead to heterogeneous mechanics and ventilation, such as ARDS and VILI.

### 4.2 Altered surface tension caused substantial alterations in ventilation dynamics

Diseases such as respiratory distress syndrome (RDS) and ARDS can lead to significant dysfunction of the pulmonary surfactant. This surfactant dysfunction can lead to substantial alteration in the dynamic surface tension during respiration, which can significantly affect the ventilation behavior. Our results indicated that altered surface tension caused substantial changes to the qualitative behavior of the P-V loop as well as the maximum volume at the same distending air pressure. There was a 20.5% reduction in maximum volume when the surface tension of the air-to-liquid interface was increased to twice the healthy value. This indicated that surfactant dysfunction, which can occur due to insufficient production or inactivation, could significantly decrease lung volumes throughout the respiratory cycle and thus impair gas exchange. This decrease in lung volumes could necessitate the need for mechanical ventilation with potentially damaging driving pressures to overcome this stiffness. However, high driving pressure mechanical ventilation in mechanical ventilation could potentially lead to the onset and spread of injuries, resulting in ventilator-induced lung injuries.

In addition, the variation of the initial alveolar radius affected the qualitative behavior of the P-V loop and the maximum volume at the same distending air pressure. This indicated that geometric parameters such as alveolar radius play an important role in the dynamic compliance of the lungs during respiration. This behavior is consistent with the lung function observed during pulmonary emphysema, which results in increased lung compliance due to damaged alveolar walls. The increased compliance in emphysema is commonly attributed to decreased parenchymal stiffness caused by the loss of alveolar septa^42^. However, this loss of alveolar septal walls results in effectively larger alveolar radii and, consequently, reduces the surface area covered by the thin liquid film^43^. The loss of surface area covered by the liquid film is captured by our model with the increased alveolar radius, indicating that reduced contribution by the surface tension of the thin liquid film could be a significant contributor to the increased compliance in emphysema. The constitutive model developed in this study can provide an essential tool for understanding the interplay between parenchymal elasticity and the surface tension of the air-to-liquid interface regulated by the pulmonary surfactant.

### 4.3 Incorporation of pulmonary surfactant enables the accurate estimation of regional parenchymal behavior

Our biophysical model can be used to predict regional mechanical stress and strains during respiration, which can be particularly important when the behavior of the lung parenchyma is heterogeneous due to conditions such as ARDS or VILI. Such conditions can lead to stress concentrations around injured regions, such as atelectatic regions, leading to increased strain and the spread of VILI. High stresses and strains could physically damage the alveolar epithelium and endothelium, resulting in altered cellular-level behavior and, consequently, organ-level dysfunction^44^. In addition, mechanical stresses modulate cellular functions in the lung, and alterations in these mechanical stresses can lead to remodeling events^1, 2^. However, the mechanotransduction behavior depends on the stresses in the alveolar septal walls, i.e., the regional stresses experienced by the lung parenchyma. Hence, it is crucial to distinguish the contribution of the lung parenchyma and pulmonary surfactant to the effective bulk modulus of the lungs. The accurate deconvolution of contributions can allow for the development of risk-stratification tools and allow clinics to be better equipped to develop clinical intervention strategies. The prediction of regional stress can also be used as an indicator of alveolar damage. Sustained increases in regional mechanical stresses can be used as an advanced biomarker to identify regions at increased risk of injury, while high levels of deformation serve as a marker for volutrauma. With the addition of recruitment dynamics in future work, the model presented in our study can be used to simulate various patient-specific ventilation protocols to reduce ventilator-induced damage.

### 4.4 Effect of natural respiration vs. mechanical ventilation on in-silico simulations

In our current study, we have investigated the effect of pulmonary surfactant on the P-V curve during positive pressure mechanical ventilation. Our novel constitutive model can also be applied to natural respiration by redefining the boundary conditions. Natural respiration occurs due to the contraction of the diaphragm and, in some cases, the intercostal muscles to expand the thoracic cavity to reduce the pressure in the pleural space. This pulls outward on the pleural surface, leading to expansion of the lungs and a decrease in alveolar pressure below atmospheric pressure. Thus, atmospheric air flows down the pressure gradient and into the parenchyma. Natural and mechanical ventilation can result in significantly different regional strains^45^ and can alter the potential for injuries. This can be simulated by applying pressure- or displacement-based boundary conditions on the outer surface of the lung model. However, accurate simulations will require either pleural pressure measurements or imaging-based displacements of the lung surface since the pleural pressure and motion of the lung surface are complex and non-uniform. Future studies will simulate natural respiration based on static CT images obtained at both the end-inspiration and end-expiration time points. Through these two images, the motion of the lung surface can be estimated based on the method reported in our previous work^46, 47^. In addition, applying image-based boundary conditions can allow us to accurately estimate stress distribution in the parenchymal tissue across the lungs.

### 4.5 Limitations

The primary limitation of the model presented in this study is the difficulty in measuring the dynamic surface tension of the air-to-liquid interface in vivo. There remains a challenge in estimating surface tension variation during respiration in vivo due to the lack of pulmonary surfactant regulation ex vivo. One possible method of assessing the effect of surfactant regulation is through inverse computational modeling. Through the measurement of the mechanical behavior of parenchymal tissue through ex-vivo testing, the surfactant behavior can be estimated by matching the pressure-volume loop predicted by the model to experimental pressure-volume measurements obtained through spirometry, plethysmography, or in ventilated subjects. Future studies will estimate surface tension through inverse modeling in diseases such as ARDS and VILI. While this method of inverse modeling would assume uniform surfactant behavior across both lungs, regional variation could be accounted for by matching model predictions to regional ventilation measured through imaging methods such as functional magnetic resonance imaging (with hyperpolarized gases)^48, 49^. Another limitation of this study is the assumption that the parenchymal tissue is completely elastic. Studies have reported that lung parenchyma demonstrates viscoelastic behavior^10, 11, 50^. In addition, the air diffusion into the parenchymal tissue was solved quasi-static, resulting in constant uniform pressure in each loading increment. Future studies will incorporate viscoelasticity into the constitutive model of the parenchymal tissue and dynamic in-silico simulations will be performed to investigate the effect of parenchymal viscoelasticity during respiration.

## 5 Conclusions

In this study, we developed a novel biophysical poroelastic model of lung parenchyma that incorporates pulmonary surfactant dynamics. We used this model to investigate the effect of altered surface tension in the air-to-liquid interface to simulate pulmonary surfactant dysfunction. The model could be an essential step towards a patient-specific model, used to investigate various lung injuries and diseases.

## 7 Acknowledgements

This work was supported by the National Institutes of Health (R00HL138288 and R56HL172052 to R.A. and R01HL151630 to B.J.S.)

## 8 Data availability

The data that support the findings of this study are available from the corresponding author, R.A., upon request.

